# PFM: perturbed flow matching for structure-based drug design

**DOI:** 10.64898/2026.07.11.737913

**Authors:** Yankai Yu, Guikun Xu, Zhuyang Xie, Yan Yang, Yongquan Jiang, Xiaobo Zhou, Kang Li

**Affiliations:** School of Computing and Artificial Intelligence, Southwest Jiaotong University, Chengdu, China; School of Artificial Intelligence, Shanghai Jiao Tong University, Shanghai, China; College of Information Engineering, Sichuan Agricultural University, Ya’an, China; Center for Computational Systems Medicine, McWilliams School of Biomedical Informatics, The University of Texas Health Science Center at Houston, Houston, TX, USA; West China Biomedical Big Data Center, West China Hospital, Sichuan University, Chengdu, China; Med-X Center for Informatics, Sichuan University, Chengdu, China

## Abstract

Generating 3D molecules that bind to specific protein targets via generative models has shown great promise in structure-based drug design. Recently, diffusion-based methods have achieved promising results, but their reliance on high sampling steps poses risks of slowing the drug discovery process due to increased time and computational costs. In this work, we propose a novel method named Perturbed Flow Matching (**PFM**), which significantly reduces sampling steps by leveraging a Flow Matching framework. PFM introduces a unique *perturbed conditional probability path design* that incorporates pocket binding site information and atom type-coordinate coupled information to enhance molecular generation performance. Experiments on CrossDocked2020 dataset demonstrate that PFM generates molecules with competitive 3D structures and state-of-the-art (SOTA) binding affinities towards the protein targets, achieving an Avg. of **-7.12**. Additionally, PFM accelerates the generation of valid molecules by a factor of **21.3**, while demonstrating potential for further improvement. The code is available at https://github.com/kurisu92725/PFM.

## 1 Introduction

Structure-based drug design (SBDD) [1] is one of the key approaches in modern drug discovery. It aims to design 3D ligand molecules with specific properties, such as high binding affinity, conditioned on a target binding site. With the development of deep learning, many generative methods have emerged to address this task. Among these, autoregressive models [2, 3, 4], which iteratively construct molecules by adding atoms and bonds, have made notable advancements. Although [5] uses fragment-based strategies to generate higher-quality molecules, this kind of method is still constrained by generation order. Recently, diffusion models [6, 7, 8] based on stochastic dynamics, have demonstrated remarkable performance. The main idea of these methods lies in denoising by inferring the reverse of the diffusion process through models. Guan et al.[9] introduced diffusion models into SBDD, and subsequent improvements have largely focused on incorporating prior knowledge to enhance the quality of the generated molecules [10, 11].

In SBDD tasks, a common expectation is to generate higher-quality molecules more rapidly. However, popular diffusion models typically require a large number of sampling steps to produce good enough results [9, 11], and fewer sampling steps pose a risk of low-quality generation [11]. To address this issue, we explore the use of Flow Matching (FM) [12, 13, 14] framework, which learns deterministic dynamics and generally offers faster sampling speeds by reducing steps [15]. However, a straight-forward application leads to poor results. Despite employing different conditional probability paths that ultimately yield the same marginal distribution both at *t* = 0 and *t* = 1, Flow Matching models produce different optimization results due to variations in paths [13, 16]. Furthermore, while the independent conditional probability path design for each variable (dimension) can form an unconditional target path through coupling [17], inspired by IPDiff [11] and EquiFM [18], we emphasize that differentiated conditional probability path designs can improve the quality of the generation.

Based on the above considerations, we propose the Perturbed Flow Matching (**PFM**) method for SBDD tasks. The perturbed conditional probability path designed in PFM maintains a correct approximation of the marginal distributions while incorporating pocket binding site information and atom type-coordinate coupled information to enhance molecular generation performance. Additionally, based on the research of simulation-free score and flow matching [16], we extend our method to incorporate stochastic dynamics without retraining.

The main contributions of this work are as follows:

1. We introduce the Perturbed Flow Matching (PFM) for molecular generation, conditioned on a target binding site, and extend it to stochastic dynamics version without retraining.
2. We propose a Full-atom Multi-stage Equivariant Network to enable flexible training of PFM.
3. Our method achieves SOTA or competitive results across most evaluation metrics, with a valid molecule generation speed 21.3 times faster than the previous SOTA method.

## 2 Related Work

### Structure-Based Drug Design

SBDD aims to generate 3D molecules with desirable properties (e.g., high affinity) conditioned on a target binding site. Skalic et al.[19] and Xu et al. [20] proposed methods for generating SMILES of molecules from given proteins. Luo et al.[2], Liu et al.[3], and Peng et al.[4] developed approaches for generating molecules in 3D Euclidean space using autoregressive techniques. Recently, the development of diffusion models has provided new solutions for SBDD tasks. Guan et al.[9], Lin et al.[21], and Schneuing et al.[22] decomposed the molecular diffusion process into continuous diffusion of atom coordinates and discrete diffusion of atom types. After generating coordinates and types, they used tools [23] to restore bonds and reconstruct the complete molecular structure. A key distinction between [9] and [21] lies in their variance schedules. Schneuing et al.[22] incorporates additional considerations such as off-the-shelf property optimization, explicit negative design, and partial molecular design with inpainting. Guan et al.[10] and Huang et al.[11] take prior knowledge into account. The former utilizes AlphaSpace2 [24] and BRICS [25] to extract the target protein subpockets and decompose molecules into fragments, while the latter introduces prior perturbations to distinguish differences among pocket binding sites by incorporating embeddings from models trained for binding affinity prediction on external datasets [26]. These diffusion-based methods typically require a large number of sampling steps to achieve high-quality generation, posing a risk of slowing down drug discovery. To address this, we propose a Flow Matching framework for SBDD to significantly enhance generation efficiency.

### Flow Matching and Applications

Flow-based generative models [27, 28, 29] faced training challenges due to the computational expense of integrating Ordinary Differential Equations (ODEs) in simulation-based training objectives. Recent advancements in simulation-free Flow Matching [12] methods have overcome this limitation. Current FM approaches focus on exploring probability paths [13, 16] and extending the theory to handle general manifolds [14]. Moreover, Discrete Flow Matching [17, 30] has emerged, enabling the generation of complex data. In the field of molecular generation, Song et al.[18] proposed an equivariant flow matching method for unconditional molecular generation, and Nori & Jin[31] applied FM methods to RNA generation. Campbell et al.[17] developed a Discrete Flow Matching approach based on Continuous Time Markov Chains (CTMCs) for protein sequence and structure generation, and Li et al.[32]’s continuous processing of amino acid types has shown promising results. The straightforward conditional probability paths in the above methods limit the performance of generated molecules in SBDD. In response, we propose a perturbed conditional probability path that incorporates pocket binding site information and atom type-coordinate coupled information, enabling differentiation of conditional probability flows. This approach enhances the quality of molecule generation for given binding sites.

## 3 Method

In this section, we provide a detailed introduction to PFM and PFM-SDE. In Sec.3.1, we introduce the standard forms of Score and Flow Matching. Sec.3.2 defines the problem and describes the construction of the basic conditional probability path. Sec.3.3 explains how to perturb basic conditional probability path. In Sec.3.4, we present a unified loss function for both deterministic and stochastic processes, and the method for obtaining perturbations in practice. In Sec.3.5, a Full-atom Multi-stage Equivariant Network is designed for training and inference.

### 3.1 Preliminaries on Score and Flow Matching

Consider a predefined marginal probability path (or target flow) *p*_*t*_ :[0, 1] × ℝ^*d*^ → ℝ^+^, which interpolates between the prior distribution *p*_0_ and the data distribution *p*_1_ = *q*. The flow *p*_*t*_ is generated by a smooth vector field 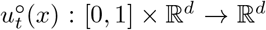, where 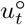 defines the deterministic process 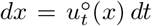 (with *continuity equation* 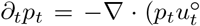) holding in this case [33]). Approximating 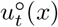 using a parameterized vector field *v*_*θ*_(*t, x*) enables the learning of a flow that matches *p*_*t*_, while the the loss function 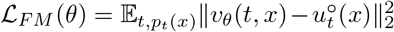 is difficult to compute due to the intractability of 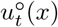. However, the availability of a conditional vector field 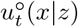 allows for optimizing an alternative objective 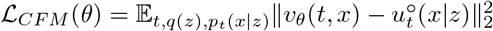, and it has been proven that ∇_*θ*_ℒ_*FM*_ (*θ*) = ∇_*θ*_ℒ_*CFM*_ (*θ*) [12, 13].

From a stochastic perspective, the marginal distribution *p*_*t*_ can evolve from *p*_0_ according to a stochastic differential equation (SDE) of the form *dx* = *u*_*t*_(*x*) *dt* + *g*(*t*) *dw*_*t*_, with the corresponding *Fokker-Planck equation* 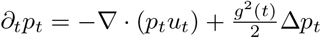 holding, where *dw*_*t*_ is a Brownian motion and Δ(*p*_*t*_ = ∇ · (∇*p*_*t*_). In this case, one can construct an associated conditional probability flow ODE: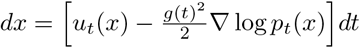, where the term in brackets is denoted as 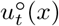; thus, the drift term can be recovered via 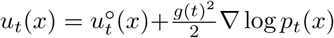. Similar to the previous case, we consider the loss functions 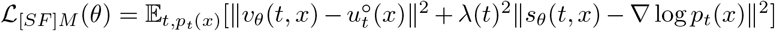 and 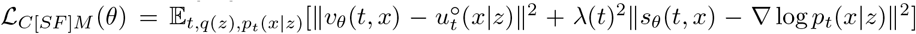. Since ∇_*θ*_ℒ_*[SF*]*M*_ (*θ*) = ∇_*θ*_ℒ_*C*[*SF*]*M*_ (*θ*), it suffices to optimize ℒ_*C*[*SF*]*M*_ (*θ*) [16].

### 3.2 Problem Definition and Basic Path Preparation

#### Problem Definition

Using generative models for SBDD involves generating ligand molecules conditioned on a given protein binding site. The protein and ligand molecule can be represented as 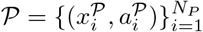 and 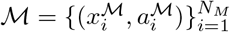, where *N*_*P*_ and *N*_*M*_ refer to the number of atoms in the protein and ligand molecule, respectively. *x*_*i*_ ∈ ℝ^3^ and *a*_*i*_ ∈ {1, 2, …, *D*} represent coordinate and types of the atom, respectively. For simplicity, we denote molecules as 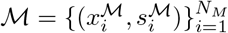, where *s*_*i*_ ∈ ℝ^*D*^ is the continuous representation of *a*_*i*_ and *D* represents the number of atom types in the ligand molecule. Furthermore, since types and coordinates of atoms in the protein remain constant during the process, we only simplify the representation of the molecule as **M** = [*X, S*], where 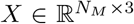 and 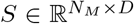. The task is then framed as modeling the conditional distribution *q*(**M**|P), with the number of atoms in the ligand molecule sampled from the prior distribution [9].

#### Continuous Processing of Atom Types

To map discrete type *a*_*i*_ to a continuous space, we employ a soft one-hot encoding operation `logit`(*a*_*i*_) = *s*_*i*_ ∈ ℝ^*D*^, with a constant value *K* > 0, and the *j*-th value of *s*_*i*_ is represented as:

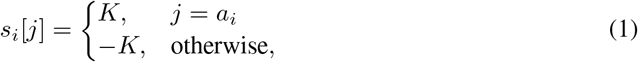

where *s*_*i*_ can be interpreted as the logit of probabilities [34].

#### Basic Conditional Probability Path Preparation

Before performing path perturbation, we select the basic paths for types and coordinates of atoms. Here, for simplicity, we adopt two similar paths for *S* and *X*:

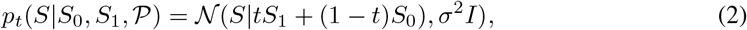

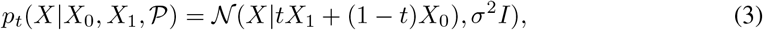

where *q*(*S*_0_) = *N*(0, *K*^2^*I*), *q*(*X*_0_) = *N*(0, *I*) and subscripts on *S* and *X* denote time. Further details on atom type continuousization and sampling can be found in Appendix A.2.

### 3.3 Perturbed Conditional Probability Path Design

Different choices of conditional probability paths *p*_*t*_(*x|z*) and conditioning distributions *q*(*z*) can lead to distinct optimization results [13, 18, 35]. Here, we introduce a novel path that does not rely on specific optimization objectives. Specifically, we propose perturbing simple conditional probability paths by incorporating pocket binding site information and atom type-coordinate coupled information to construct a new conditional probability path, which we term a “perturbed conditional probability path”. The form is as follows:

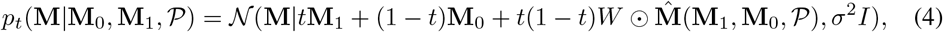

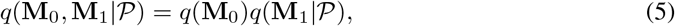

where 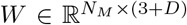 represents the weight matrix. 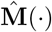 is a function, which, together with *W*, is required to ensure that *p*_*t*_(**M**|**M**_0_, **M**_1_, *P*) can be integrated over **M**_0_ and **M**_1_ to obtain the marginal probability *p*_*t*_(**M**|*P*). ⊙ denotes the Hadamard product. In the independent coupling represented by Eq.5, translational invariance in coordinate *X*, where *p*(*X* + **t**) = *p*(*X*) and **t** denotes the translation, can lead to non-normalizable probability distributions. To address this issue, we use the centroid of the pocket binding site as the origin [36]. In practice, we further employ the following simplified perturbed conditional probability path, as illustrated in Fig.1:

**Figure 1:**
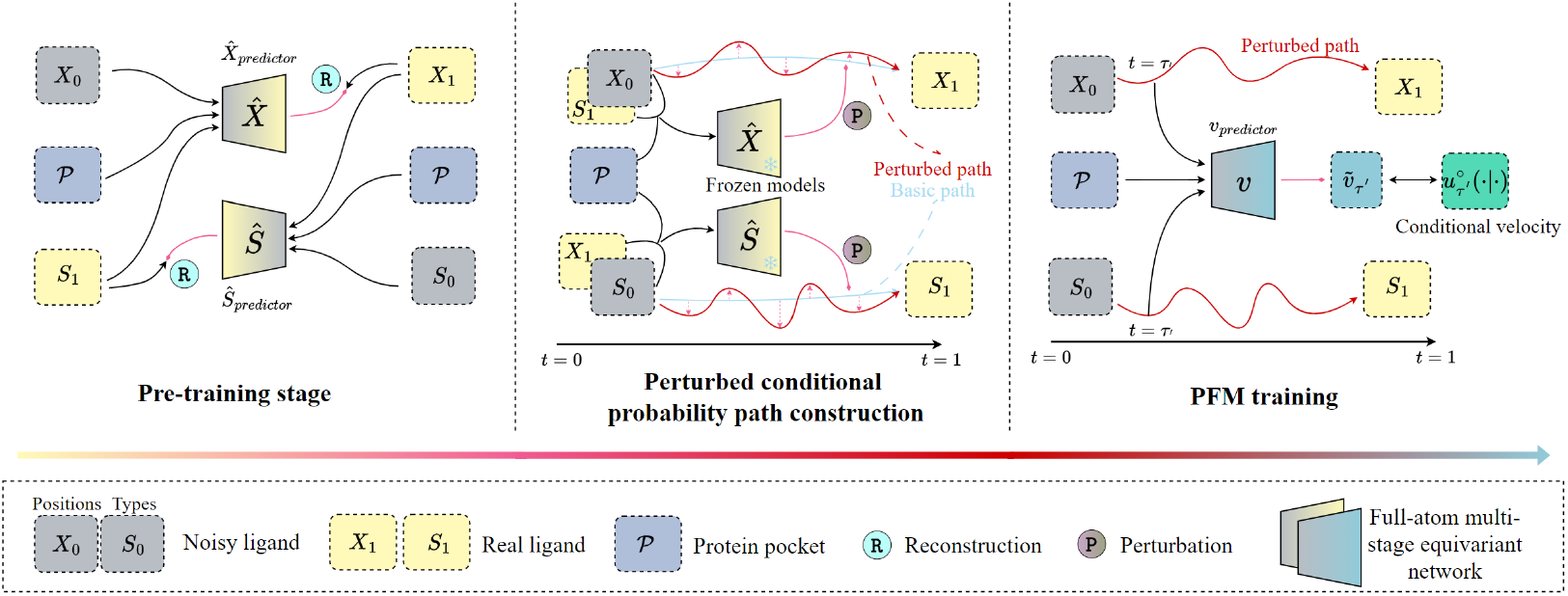
Specific training procedure of PFM. The **middle** is the crucial construction of the perturbed conditional probability path.

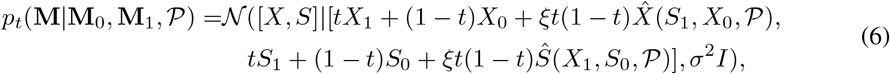

where 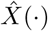 and *Ŝ*(·) are functions, with 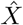 and *Ŝ* in the following referring to the perturbations from computation. In practice, we find that using neural networks to generate these perturbations works well. The magnitude of the perturbation is jointly controlled by *ξ* and *t*(1 − *t*).

Eq.4 and 6 describe the conditional flow of each element in the matrix **M** or [*X, S*]. The non-“straight” nature of the perturbed conditional probability paths reflected in Eq.6 contributes to the distinguishability between different trajectories, which may offer potential benefits for generative model construction [11]. Moreover, based on the results in [13, 18, 35], we would like to emphasize that *<u>“straight” paths do not always lead to good generative performance</u>*. As demonstrated in the experimental results of Sec.4, proposed perturbed paths improve molecular generation performance.

The details of 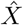 and *Ŝ* involved are introduced at the end of Sec.3.4. According to Sec.3.1 and **Theorem 3** of [12], the conditional vector field can be expressed as follows:

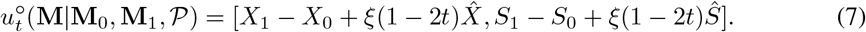

Notably, *p*_*t*_(**M**|**M**_0_, **M**_1_, *P*) can be efficiently sampled, and 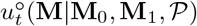 can be easily computed, which makes the optimization of *θ* on ℒ_*CFM*_ feasible. As *σ* → 0, the marginal boundary probabilities converge to *q*(**M**_0_) and *q*(**M**_1_|P) according to **Proposition 3.3** in [13].

When *p*_*t*_ is generated by the stochastic process described in Sec.3.1, the conditional score is given by:

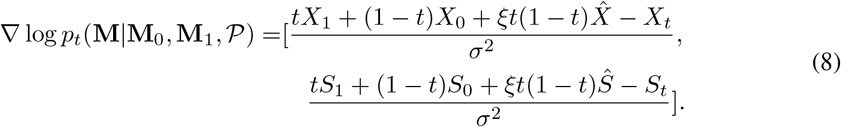

Further details on constructing flows for the SBDD task can be found in Appendix A.

### 3.4 Training Objective

#### Score and Flow Matching Loss

Directly predicting drift and score function requires training separate networks for the deterministic and stochastic processes [16]. Inspired by [7], we derive a unified loss function for both processes through reparameterization and setting coefficients to unity:

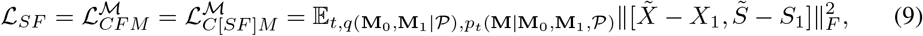

where *t* ∼ *U*(0, 1), and 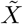 and *Ŝ* are predictions from the target neural network, distinct from the networks previously used to generate perturbations.This equation implies that we only need to train the deterministic process, while enabling flexible sampling methods for both processes. The detailed reparameterization is provided in Appendix A.1. However, *L*_*SF*_ focuses solely on individual atoms, lacking consideration for local molecular structures and interactions with the protein. Here, we propose incorporating two simple auxiliary losses to account for these considerations.

#### Local Loss

From an intramolecular perspective, it is important to incorporate insights into local structure, such as bond lengths, bond angles, and torsion angles. For simplicity, we introduce only the bond length loss:

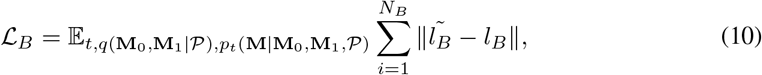

where *N*_*B*_ denotes the number of bonds in a ligand molecule, and *l*_*B*_ is ground truth of bond length.

#### Clash Loss

From the perspective of molecule–pocket interactions, steric clashes are a critical consideration. Therefore, we introduce a clash loss to penalize atoms that intrude into the protein interior. Following [37, 10], we choose {*x* ∈ ℝ^3^ : *S*(*x*) = *γ*} where 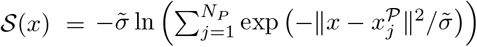 as the descriptor of the protein surface. Subsequently, we define a clash loss as:

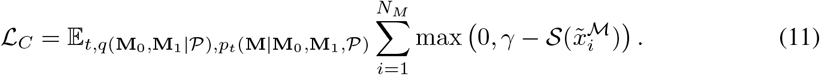

The overall loss function for PFM or PFM-SDE is thus expressed as:

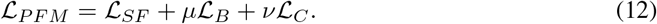

#### Perturbation Network and Reconstruction Loss

To enhance the stability of PFM training and reduce memory usage, we leverage pre-trained predictors for the perturbations 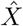 and *Ŝ* as shown in Fig.1. Specifically, we introduce a molecular reconstruction task given protein pocket *P* and atom types *S* or coordinates *X* of ligand molecule. Their corresponding loss functions are defined as:

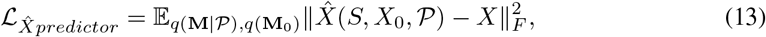

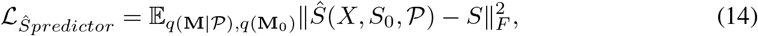

where **M** = [*X, S*]. The pre-trained predictors introduce meaningful and reasonable perturbations into basic probability paths. However, this training strategy raises concerns about the effectiveness of this kind of perturbation, such as the difference between paths with mean 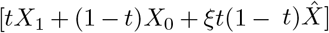 and [*tX*_1_ + (1 − *t*)*X*_0_ + *ξt*(1 − *t*)*X*_1_]. These concerns will be addressed in Sec.4.3.

### 3.5 Full-atom Multi-stage Equivariant Network

Inspired by recent advancements in equivariant neural networks [38, 39, 40] and protein-ligand binding affinity prediction [41], a multi-stage approach is proposed to update features and positions.

First, three atom-level *k*-nearest neighbor (knn) graphs are constructed: *G*_*P*_, *G*_*M*_, and *G*_*I*_ representing the protein, ligand molecule, and protein-ligand interaction graphs, respectively. The value of *k* differs for each graph. After embedding the initial node features into the same dimension for both protein and molecule atoms, the protein atom features are updated as follows in each layer:

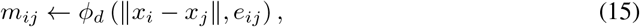

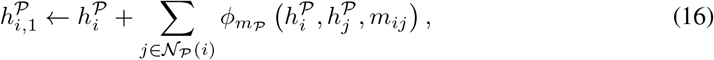

where 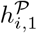 represents the updated feature of node *i* in graph *G*_*P*_, *N*_*P*_ (*i*) denotes the set of neighboring nodes’ indices of node *i* in *G*_*P*_, *e*_*ij*_ represents edge *ij* in related *G*, and *m*_*ij*_ denotes message. Since the network ultimately obtains atom representations in ligand molecules, the updating strategy in *G*_*M*_ differs slightly from that in *G*_*P*_. **In the first stage**, atomic interactions within the ligand molecule are considered, and the type features and positions are updated as follows:

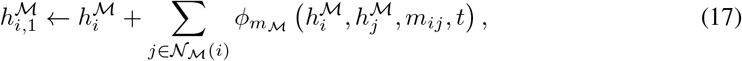

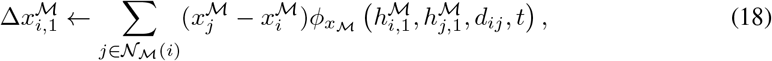

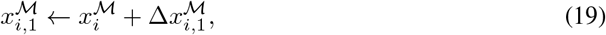

where *d*_*ij*_ is the Euclidean distance between nodes *i* and *j*.

**In the second stage**, atomic interactions between the ligand molecule and the protein are considered, and the type features and positions are updated as follows:

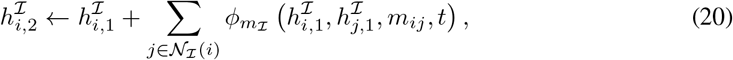

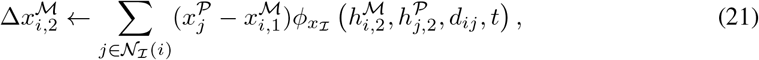

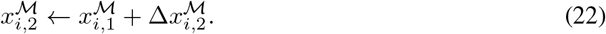

Here, Eq.21 distinguishes the atomic origin. During both stages, the subscripts on *h* and *x* (e.g., 1, 2) denote updates within the same layer. *m*_*ij*_ and *d*_*ij*_ change dynamically due to updating.

After *L* layers of updates, the network outputs 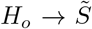 and 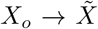, which are then passed to the loss function ℒ_*PFM*_ for parameter optimization. The same network structure is adopted for pre-trained predictors. Further details on pretraining and PFM training can be found in Appendix C, while procedures for training and sampling are provided in Algorithm 1 and Algorithm 2.

## 4 Experiments

### 4.1 Experimental Setup

#### Dataset

Following the previous work [2, 4], we train and evaluate PFM on CrossDocked2020 dataset [42]. The data preprocessing and splitting methods used by [9] and [2] are adopted, refining the 22.5 million docked binding complexes to retain high-quality docking poses (RMSD between the docked pose and the ground truth < 1 Å) and diverse proteins (sequence identity 30%). Ultimately, 99,900 qualified complexes are utilized for training, and 100 novel proteins are selected for testing.

#### Baseline

In this study, the proposed method is compared with several recent representative approaches: **LiGAN** [43] is a CVAE model implemented on atomic density grids. **AR** [2], **Pocket2Mol** [4] and **GraphBP** [3] generate molecules autoregressively by leveraging contextual information. **TargetDiff** [9], **DecomposeDiff** [10], and **IPDiff** [11] are diffusion-based models designed for molecular generation.

#### Evaluation

The generated molecules are evaluated from two perspectives: **Target Binding Results** and **Molecular Conformations and Properties**.

- **Target Binding Results** The mean and median of *Vina Score, Vina Min, Vina Dock*, and *High Affinity* are used to evaluate binding affinities [2, 43]. *Vina Score* directly estimates the binding affinity of generated molecules to the target, *Vina Min* performs local minimization before estimation, *Vina Dock* involves a re-docking process reflecting the best possible binding affinity, and *High Affinity* indicates the percentage of molecules generated for each test protein that bind better than the reference molecules. In addition, *Clash Ratio* and *RMSD* are considered to evaluate the geometry of docking poses. Specifically, *CCA*. denotes the ratio of atoms that clash with protein atoms to the total number of atoms, and *CM*. indicates the ratio of molecules with clashes [44]. *%*<*2Å* denotes the percentage of generated molecules with RMSD less than 2Å from the redocked poses, serving as a measure of the model’s ability to learn the docking poses [45, 46].
- **Molecular Conformation and Properties** Molecular conformation is evaluated by measuring the Jensen-Shannon divergences (JSD) between the bond distance distributions of reference and generated molecules. The strain energy (*SE*) of the generated molecules is also considered, reflecting the energetic stability of conformation [47]. For molecular properties, *QED, SA, Diversity, Similarity, Validity*, and *Uniqueness* are used [2, 43, 48, 49]. *QED* quantifies the drug-likeness of molecules, *SA* evaluates their synthetic accessibility, and *Diversity* represents the average pairwise dissimilarity between the molecules generated for a given pocket binding site. *Similarity* measures the average similarity between generated molecules and their references. *Validity* refers to the proportion of generated molecules that can be reconstructed by RDKit and have binding affinities <0. *Uniqueness* indicates the proportion of valid molecules with different canonical SMILES.

### 4.2 Main Results

#### Target Binding Results

As shown in Tab.1, for **binding affinity**, PFM significantly outperforms the autoregressive model Pocket2Mol, with notable improvements of 38.5%, 18.1%, and 16.4% in the average Vina Score, Vina Min, and Vina Dock, respectively. Compared with IPDiff, PFM achieves competitive results without introducing additional prior knowledge (e.g., binding affinity prediction tasks), showing a 10.9% improvement in average Vina Score, a slight increase of 1.7% in average Vina Min, and a minor decrease of 2.9% in average Vina Dock. It is important to note that, as described in Sec.4.1, Vina Score best reflects the model’s molecular generation and docking ability. In addition, larger generated molecules or distortions introduced by the model could potentially lead to better Vina Scores. However, as shown by the average number of atoms in Tab.1 and the strain energy results in Tab.3, PFM does not appear to benefit significantly from either of these factors, which could otherwise compromise the fairness of comparisons or the stability of generated molecules. For **geometry evaluation**, PFM outperforms other diffusion-based methods in Clash Ratio and RMSD. LiGAN achieves the best clash ratio, primarily because its grid-based modeling effectively avoids most steric clashes.

**Table 1:** Summary of target binding results of reference molecules and molecules generated by our model and other non-diffusion (**Others**) and diffusion-based (**Diffusion**) baselines. (↑) / (↓) denotes a larger / smaller number is better. Top 3 results are highlighted with **bold text** and <u>underlined text</u>, respectively.

| Methods | | Vina Score (↓) | | Vina Min (↓) | | Vina Dock (↓) | | High Affinity (↑) | | Clash Ratio (↓) | | RMSD (↑) | # Atoms /mol. |
| --- | --- | --- | --- | --- | --- | --- | --- | --- | --- | --- | --- | --- | --- |
|  |  | Avg. | Med. | Avg. | Med. | Avg. | Med. | Avg. | Med. | CCA. | CM. | %<2 Å | Avg. |
| Reference |  | -6.36 | -6.46 | -6.71 | -6.49 | -7.45 | -7.26 | - | - | - | - | 34.0 | 22.8 |
| **Others** | liGAN | - | - | - | - | -6.33 | -6.20 | 21.1% | 11.1% | **0.0096** | **0.0718** | **43.8** | 19.7 |
|  | GraphBP | - | - | - | - | -4.80 | -4.70 | 14.2% | 6.7% | 0.8634 | 0.9974 | - | 23.1 |
|  | AR | -5.75 | -5.64 | -6.18 | -5.88 | -6.75 | -6.62 | 37.9% | 31.0% | - | - | - | - |
|  | Pocket2Mol | -5.14 | -4.70 | -6.42 | -5.82 | -7.15 | -6.79 | 48.4% | 51.0% | 0.0576 | 0.4499 | 30.8 | 17.7 |
| **Diffusion** | TargetDiff | -5.47 | -6.30 | -6.64 | -6.83 | -7.80 | -7.91 | 58.1% | 59.1% | 0.0483 | 0.4920 | 29.4 | 24.2 |
|  | DecompDiff | -5.67 | -6.04 | -7.04 | -7.09 | -8.39 | -8.43 | 64.4% | 71.0% | 0.0462 | 0.5248 | 23.9 | 20.9 |
|  | IPDiff | -6.42 | -7.01 | -7.45 | **-7.48** | **-8.57** | **-8.51** | 69.5% | 75.5% | 0.0313 | 0.3463 | - | 24.5 |
| **FM** | **PFM** | **-7.12** | **-7.18** | **-7.58** | **-7.32** | **-8.32** | **-8.26** | **72.1%** | **76.5%** | **0.0145** | **0.2040** | **36.7** | 22.6 |
|  | **PFM-SDE** | <u>-6.99</u> | <u>-7.13</u> | <u>-7.48</u> | <u>-7.32</u> | -8.29 | -8.23 | <u>71.2%</u> | **76.6%** | <u>0.0155</u> | <u>0.2131</u> | <u>37.3</u> | 22.5 |

#### Molecular Conformation and Properties

JSD values between bond distance distributions of reference and generated molecules are calculated and summarized in Tab.2. The results demonstrate that molecules generated by PFM-SDE achieved SOTA performance in five out of eight common bond types. While PFM performs slightly below PFM-SDE, it delivers competitive results compared with IPDIFF (the previous SOTA). For SE in Tab.3, PFM outperforms diffusion-based methods, indicating that it generates more stable docking conformations. Regarding **molecular properties**, PFM achieves superior or competitive performance across most metrics compared with diffusion-based methods, though it performs significantly worse on SA. However, as shown in the ablation studies in Tab.45, this deficiency primarily arises from the trade-off between QED and SA induced by the perturbed conditional probability path and the ℒ_*B*_ term, rather than a direct decline in SA. Nonetheless, according to [50], the SA scores of molecules generated by PFM remain within an acceptable range and do not hinder the rough screening of drug discovery [9, 10].

**Table 2:** Jensen-Shannon divergence between bond distance distributions of the reference molecules and the generated molecules, and lower values indicate better performances. “-”, “=“, and “:” represent single, double, and aromatic bonds, respectively. Top 3 results are highlighted with **bold text** and <u>underlined text</u>, respectively.

| Bond | liGAN | GraphBP | AR | Pocket2 Mol | Target Diff | Decomp Diff | IP Diff | PFM | PFM-SDE |
| --- | --- | --- | --- | --- | --- | --- | --- | --- | --- |
| C—C | 0.601 | <u>0.368</u> | 0.609 | 0.496 | <u>0.369</u> | **0.359** | 0.386 | 0.395 | 0.375 |
| C=C | 0.665 | <u>0.530</u> | 0.620 | 0.561 | <u>0.505</u> | 0.537 | <u>0.245</u> | <u>0.172</u> | **0.165** |
| C—N | 0.634 | 0.456 | 0.474 | 0.416 | 0.363 | 0.344 | <u>0.298</u> | <u>0.266</u> | **0.254** |
| C=N | 0.749 | 0.693 | 0.635 | 0.629 | 0.550 | 0.584 | <u>0.238</u> | **0.147** | **0.147** |
| C—O | 0.656 | 0.467 | 0.492 | 0.454 | 0.421 | 0.376 | <u>0.366</u> | <u>0.317</u> | **0.309** |
| C=O | 0.661 | 0.471 | 0.558 | 0.516 | 0.461 | 0.374 | **0.353** | 0.411 | <u>0.392</u> |
| C:C | 0.497 | 0.407 | 0.451 | 0.416 | 0.263 | 0.251 | 0.169 | 0.188 | **0.165** |
| C:N | 0.638 | 0.689 | 0.552 | 0.487 | 0.235 | 0.269 | **0.128** | <u>0.187</u> | <u>0.181</u> |

**Table 3:** Summary of molecular conformation and properties of reference molecules and molecules generated by our model and other baselines. (↑) / (↓) denotes a larger / smaller number is better. Top 3 results are highlighted with **bold text** and <u>underlined text</u>, respectively.

| Methods | QED (↑) | | SA (↑) | | Diversity (↑) | | SE (↓) | | | Similarity (↓) | Validity (↑) | Uniqueness (↑) |
| --- | --- | --- | --- | --- | --- | --- | --- | --- | --- | --- | --- | --- |
|  | Avg. | Med. | Avg. | Med. | Avg. | Med. | 25% | 50% | 75% | Avg. | Avg. | Avg. |
| Reference | 0.48 | 0.47 | 0.73 | 0.74 | - | - | 34 | 107 | 196 | - | 100% | - |
| liGAN | 0.39 | 0.39 | 0.59 | 0.57 | 0.66 | 0.67 | 212 | 619 | 1855 | 0.22 | - | 87.28% |
| GraphBP | 0.43 | 0.45 | 0.49 | 0.48 | **0.79** | **0.78** | **46** | **105** | 464 | **0.15** | - | **100%** |
| Pocket2Mol | **0.56** | **0.57** | **0.74** | **0.75** | 0.69 | <u>0.71</u> | <u>102</u> | <u>186</u> | **374** | 0.26 | **98.31%** | **100%** |
| TargetDiff | 0.48 | 0.48 | 0.58 | 0.58 | 0.72 | <u>0.71</u> | 369 | 1243 | 13871 | 0.30 | 90.35% | 99.63% |
| DecompDiff | 0.45 | 0.43 | <u>0.61</u> | <u>0.60</u> | 0.68 | 0.68 | 379 | 983 | 4133 | 0.34 | 71.96% | 99.99% |
| IPDiff | <u>0.52</u> | <u>0.53</u> | <u>0.61</u> | <u>0.59</u> | <u>0.74</u> | <u>0.73</u> | 600 | 2720 | 6299 | <u>0.18</u> | 88.16% | **100%** |
| **PFM** | <u>0.49</u> | <u>0.50</u> | 0.51 | 0.51 | <u>0.73</u> | <u>0.71</u> | 229 | 783 | <u>1708</u> | <u>0.20</u> | <u>95.12%</u> | **100%** |

**Table 4:** The influence of perturbed conditional probability path.

| Methods | Vina Score (↓) | | Vina Min (↓) | | QED (↑) | | SA (↑) | |
| --- | --- | --- | --- | --- | --- | --- | --- | --- |
|  | Avg. | Med. | Avg. | Med. | Avg. | Med. | Avg. | Med. |
| BFM* | -4.27 | -4.94 | -6.04 | -5.94 | 0.46 | 0.47 | 0.50 | 0.51 |
| BFM | -5.77 | -6.27 | -6.76 | -6.71 | 0.47 | 0.48 | <u>0.54</u> | <u>0.54</u> |
| BFM-Ref | <u>-6.48</u> | <u>-6.89</u> | -7.27 | -7.10 | **0.49** | **0.50** | 0.51 | 0.51 |
| LPFM | -6.72 | -6.88 | <u>-7.37</u> | <u>-7.13</u> | **0.49** | **0.50** | 0.51 | 0.50 |
| RPFM | <u>-6.86</u> | <u>-7.17</u> | <u>-7.51</u> | **-7.33** | 0.48 | 0.49 | <u>0.52</u> | <u>0.52</u> |
| PFM* | <u>-6.48</u> | -6.49 | -6.79 | -6.45 | 0.43 | 0.43 | **0.57** | **0.57** |
| PFM | **-7.12** | **-7.18** | **-7.58** | <u>-7.32</u> | **0.49** | **0.50** | 0.51 | 0.51 |

**Table 5:** The effect of different loss terms.

| $\mathcal{L}_{SF}$ | $\mathcal{L}_B$ | $\mathcal{L}_C$ | Vina Score (↓) | | Vina Min (↓) | | QED (↑) | | SA (↑) | |
| --- | --- | --- | --- | --- | --- | --- | --- | --- | --- | --- |
|  |  |  | Avg. | Med. | Avg. | Med. | Avg. | Med. | Avg. | Med. |
| ✓ | ✗ | ✗ | -6.48 | -6.49 | -6.79 | -6.45 | 0.43 | 0.43 | **0.57** | **0.57** |
| ✓ | ✓ | ✗ | -6.88 | -7.06 | -7.43 | -7.18 | 0.49 | 0.50 | 0.52 | 0.51 |
| ✓ | ✗ | ✓ | -6.45 | -6.48 | -6.83 | -6.51 | 0.42 | 0.42 | **0.57** | 0.56 |
| ✓ | ✓ | ✓ | **-7.12** | **-7.18** | **-7.58** | **-7.32** | **0.49** | **0.50** | 0.51 | 0.51 |

#### Faster Sampling and Performance Gains from Perturbation

As shown in Tab.6, PFM achieves a 21.3x speed-up in sampling, benefiting from the inherent efficiency of Flow Matching that require fewer sampling steps, as well as the flexibility of the proposed network in graph construction. However, this acceleration comes at the cost of poor baseline performance, as evidenced by the BFM* results in Tab.4. The proposed perturbed conditional probability path significantly improves docking energy and introduces a trade-off between QED and SA (as shown by PFM*), while the simple auxiliary loss further enhances model effectiveness. The potential for faster sampling can be found in Appendix D.

**Table 6:** Training and inference time.

| Methods | Training_t(hrs) | Inference_t(hrs) |
| --- | --- | --- |
| TargetDiff | 5.57 | 22.11 |
| IPDiff | 9.53 | 66.68 |
| BFM | 4.31 | 1.17 |
| $\hat{X}$ predictor | 4.22 | - |
| $\hat{S}$ predictor | 4.17 | - |
| PFM | 6.31 | 3.13 |

### 4.3 Ablation Studies

PFM introduces an FM-based molecular generation approach for SBDD, featuring a novel perturbed conditional probability path design and a well-performed loss function construction. A comprehensive ablation analysis validates the effectiveness of these components. Further results and analysis on reducing the number of sampling steps, the impact of perturbation coefficients, and the differences in predicted fields caused by perturbed conditional probability paths can be found in Appendix D.

#### Effect of Perturbed Conditional Probability Path

The perturbed conditional probability path in PFM is hypothesized to improve molecular generation performance. Tab.4 presents the impact of different path choices on the quality of generated molecules, all utilizing Gaussian paths with identical variance but differing means. Taking the conditional probability path on *X* as an example, the mean for Basic Flow Matching (**BFM**) is [*tX*_1_+(1−*t*)*X*_0_]; for **BFM-Ref**, [*tX*_1_+(1−*t*)*X*_0_ +*ξt*(1−*t*)*X*_1_]; for Left-Shifted PFM (**LPFM**), 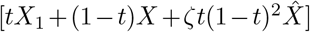; and for Right-Shifted PFM (**RPFM**), 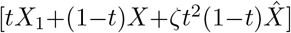. Note that **BFM\*** and **PFM\*** indicate models trained with consideration of ℒ_*SF*_ only. The results demonstrate that the proposed perturbed paths improve overall molecular generation performance (as shown by comparisons of PFM with BFM and BFM-Ref, or PFM* with BFM*) and confirm the effectiveness of PFM under different perturbation strategies (as evidenced by comparisons of LPFM, RPFM, PFM with BFM and BFM-Ref).

#### Impact of the Loss Function

The proposed loss function significantly enhances PFM’s molecular generation performance. Tab.5 presents an ablation study of the loss terms, indicating that ℒ_*B*_ contributes a lot to performance improvement. ℒ_*C*_ provides slight benefits to molecular generation performance only when combined with ℒ_*B*_ and ℒ_*SF*_.

## 5 Conclusions

In this paper, a method called PFM for the SBDD task is proposed for the first time. Building on a basic FM framework, PFM enhances molecular generation performance by designing a more differentiated perturbed path, which incorporates pocket binding site information and atom type-coordinate coupled information into a basic conditional probability path. Additionally, the introduction of a flexible full-atom multi-stage equivariant network and a simple auxiliary loss also contribute to improved results. Experiments demonstrate the efficiency of the model and the effectiveness of its components, and highlight PFM’s potential to generate competitive molecules with faster sampling.

## 6 Limitations and Future Work

PFM have been demonstrated to be effective in this paper. However, certain limitations remain. From the perspective of generative algorithms, PFM explores the construction of generative paths to a certain extent. The effectiveness of this approach lacks large-scale validation in multimodal generation tasks, due to computational constraints. Moreover, in various generative tasks, the design and coupling of modality-specific paths warrant deeper investigation. From the SBDD perspective, PFM focuses on local structural information through the loss function during generation, which may be insufficient. Additional structural constraints should be incorporated. Our future work will also explore matching of multi-level molecular generation, the integration of domain-specific priors, and the application of large models.

## A. Score and Flow Matching in the SBDD Task

### A.1 Obtaining ℒ_*CFM*_ **and** ℒ_*C*[*SF*]*M*_ through Reparameterization

According to Sec.3.1, the conditional flow matching loss in the SBDD task is defined as:

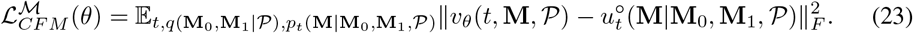

Substituting Eq.7 into Eq.23:

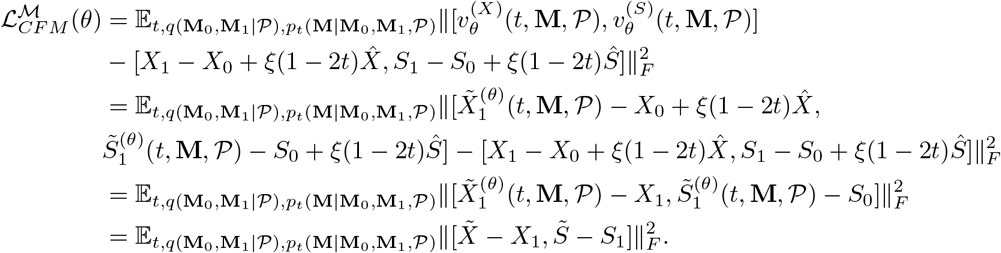

It is important to note that, taking the vector field of position *X* as an example, the perturbation term contributing to the predicted vector field in the second equality is actually 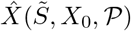. Strictly enforcing this term increases training complexity and degrades the quality of the final generation. Our simple loss function provided is sufficient to optimize the vector field prediction, and this simplification does not compromise the inference process. The results in Tab.7 validate the effectiveness of our simplification.

**Table 7:** The influence of loss function and conditioning distribution.

| Methods | Vina Score (↓) | | Vina Min (↓) | | QED (↑) | | SA (↑) | | Loss (↓) | |
| --- | --- | --- | --- | --- | --- | --- | --- | --- | --- | --- |
|  | Avg. | Med. | Avg. | Med. | Avg. | Med. | Avg. | Med. | Training | Test |
| PFM-TL | -6.83 | -6.98 | -7.49 | -7.23 | 0.47 | 0.47 | 0.50 | 0.51 | - | - |
| PFM-IC | -4.42 | -4.76 | -7.58 | -7.16 | 0.49 | 0.50 | 0.42 | 0.39 | 5.48 | 9.83 |
| PFM | -7.12 | -7.18 | -7.58 | -7.32 | 0.49 | 0.50 | 0.51 | 0.51 | 1.27 | 2.28 |

From Sec.3.1, the conditional score matching loss part of ℒ_*C*[*SF*]*M*_ in the SBDD task is defined as:

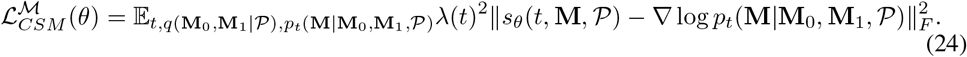

Substituting Eq.8 into Eq.24, and following a derivation similar to that of ℒ_*CFM*_ :

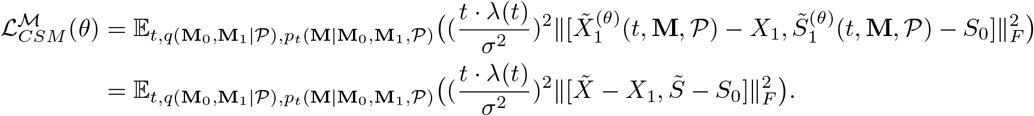

During training, we set 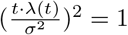 to ensure a stable training process. Now, we observe that the score matching loss and flow matching loss share the same form, and further normalization of the coefficients leads to Eq.9. This implies that we only need to train the network on the deterministic dynamics, and during inference, we can freely choose between the deterministic and stochastic dynamics from Sec.3.1 to generate molecules, without retraining.

### A.2 Flow of Atom Type on Simplex

Based on Eq.1, *s*_*i*_ undergoes a `softmax` operation, resulting in a normalized probability distribution. The *i*-th dimension of this distribution approaches 1, while the remaining dimensions converge to 0, indicating that the probability mass is correctly concentrated on the atom type *a*_*i*_. Specifically, `softmax` (*s*_*i*_) represents a point on the D-category probability simplex Δ^*D−*1^, where 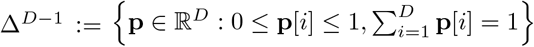 is a smooth manifold in ℝ^*D*^ [51, 52, 53]. Each point on Δ^*D−*1^ represents a categorical distribution over *D* classes. Instead of directly constructing flows on Δ^*D−*1^, as detailed in [32], we define a conditional flow in the logit space ℝ^*D*^ with a Gaussian prior, as shown in Eq.2.

During inference, Euler steps are performed to solve the probability flow in the logit space, and atom types can then be sampled from the corresponding probability vector on the simplex:

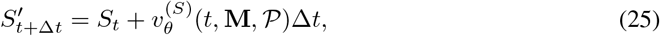

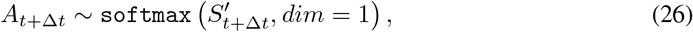

where 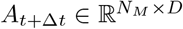 denotes atom type sampled according to Eq.26.

To improve consistency between the logit space and the simplex space during generation, the predicted atom types are mapped back to the logit space throughout the iterative process:

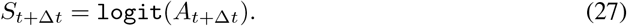

Additionally, according to Appendix A.1 of [32], a rigorous and correct approach involves reducing one degree of freedom in the logit space to ensure a one-to-one mapping between the simplex and the logit space. However, this setting is not adopted in the implementation of [32], and it does not show a significant impact in PFM.

### A.3 Modification of Independent Coupling (Eq.5)

Tong et al.[13] used mini-batch optimal transport to improve Flow Matching (OT-FM), while Klein et al.[35] extended this with Equivariant Optimal Transport Flow Matching (EOT-FM) to provide improved conditioning distributions. These approaches significantly shorten the path length during training, empirically improving both training stability and generation quality. However, applying EOT to the SBDD task poses practical challenges. Specifically, variations in atom counts among ligands within a batch make mini-batch EOT infeasible. Inspired by the explanation of **Discrete Optimal Transport** in Appendix B.1 of [35], we propose a method to shorten the path length and devise a corresponding sampling strategy. When the cost function *c*(*X*_0_, *X*_1_) = *d*(*X*_0_, *X*_1_), where *d* denotes the Euclidean distance, the sampling process is defined as:

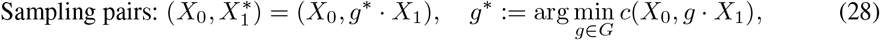

where *X*_0_ and *X*_1_ sampled according to Eq.5, and *G* is a compact (topological) group acting on an *n*-dimension Euclidean space *X* via isometries. This includes the symmetric group S_*n*_, the orthogonal group O_*n*_, and all their subgroups. Fig.2 provides a more intuitive depiction of the differences in basic conditional probability paths (flow matching differences) during training. In practice, for simplicity, only the symmetric group 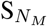 is considered, and s^*\**^ is obtained using Hungarian algorithm [54]. The impact of the modified conditioning distributions on the performance of generated molecules is detailed in Tab.7.

**Figure 2:**
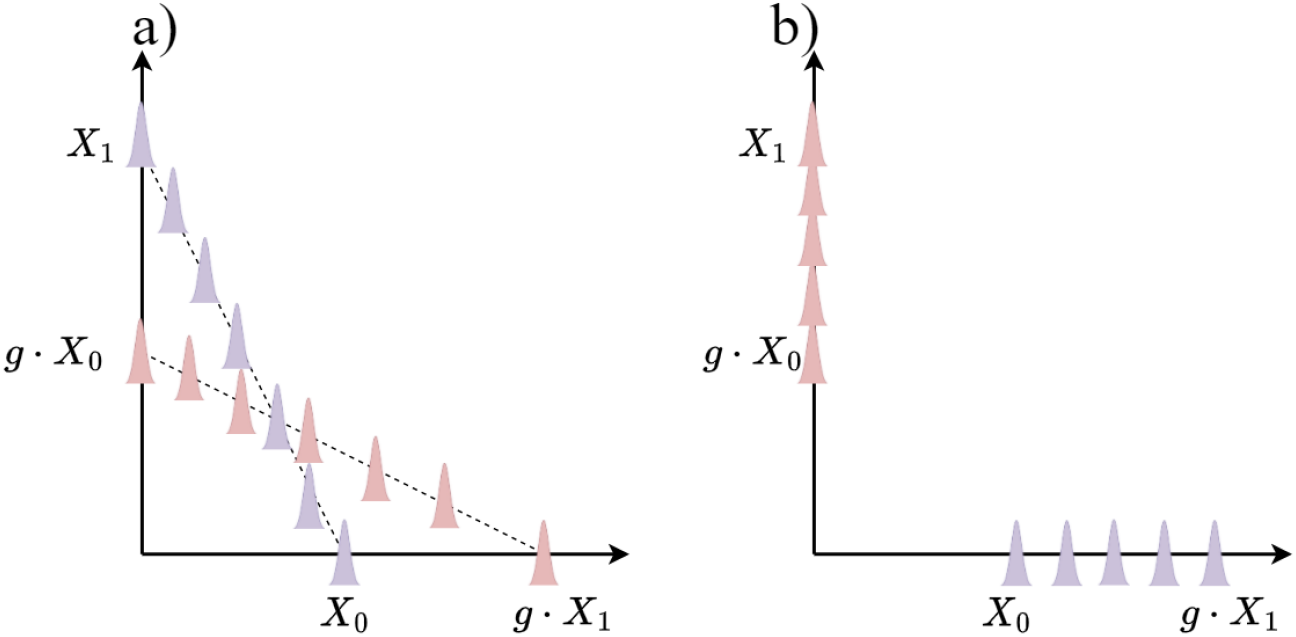
Different sampling methods lead to distinct basic conditional probability paths during training: a) the original path and b) the modified path.

## B Training and Sampling Procedure

The training and sampling procedures are summarized as Algorithm 1 and Algorithm 2.

### Algorithm 1 Training Procedure of PFM

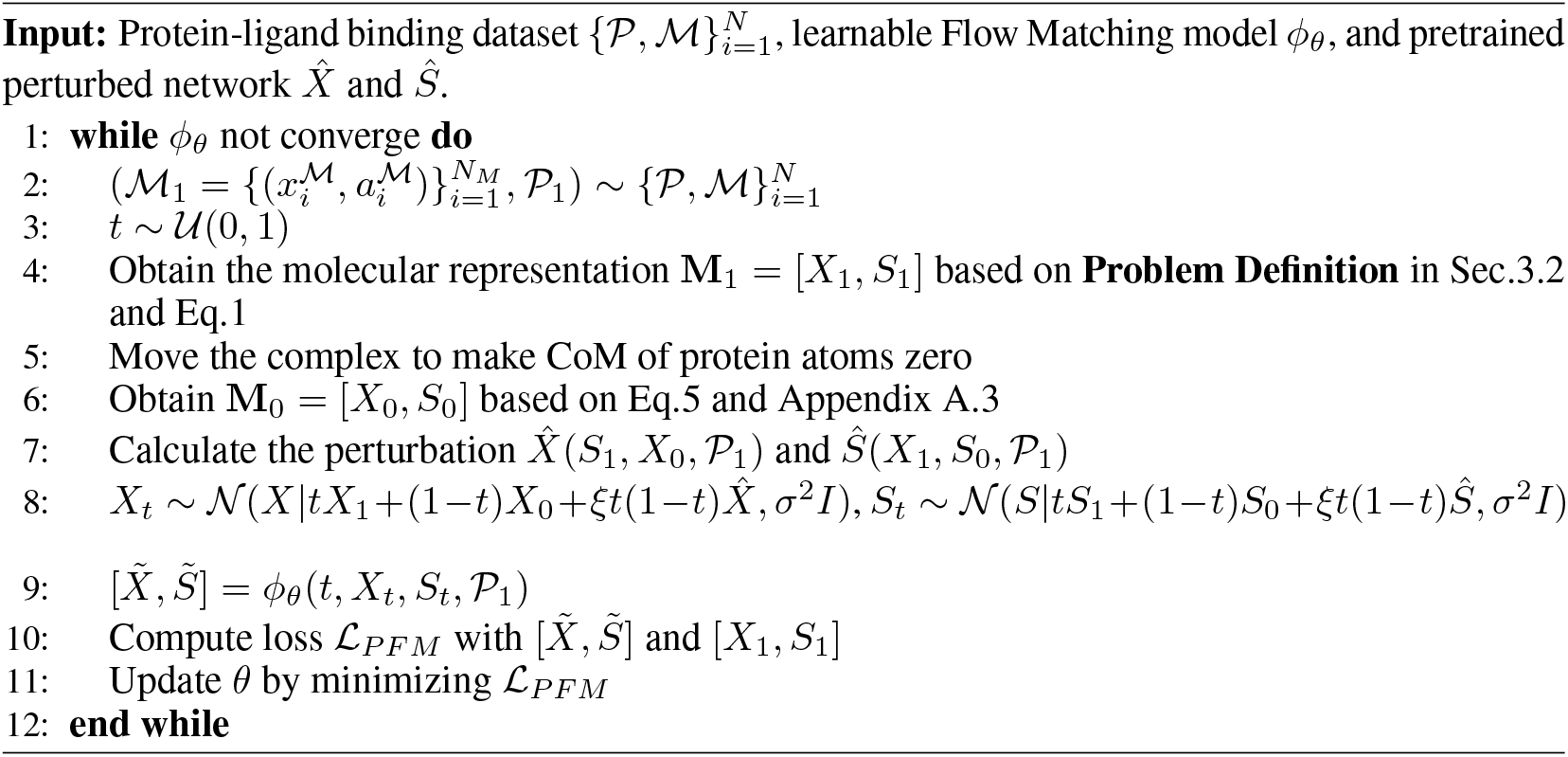

### Algorithm 2 Sampling Procedure of PFM and PFM-SDE

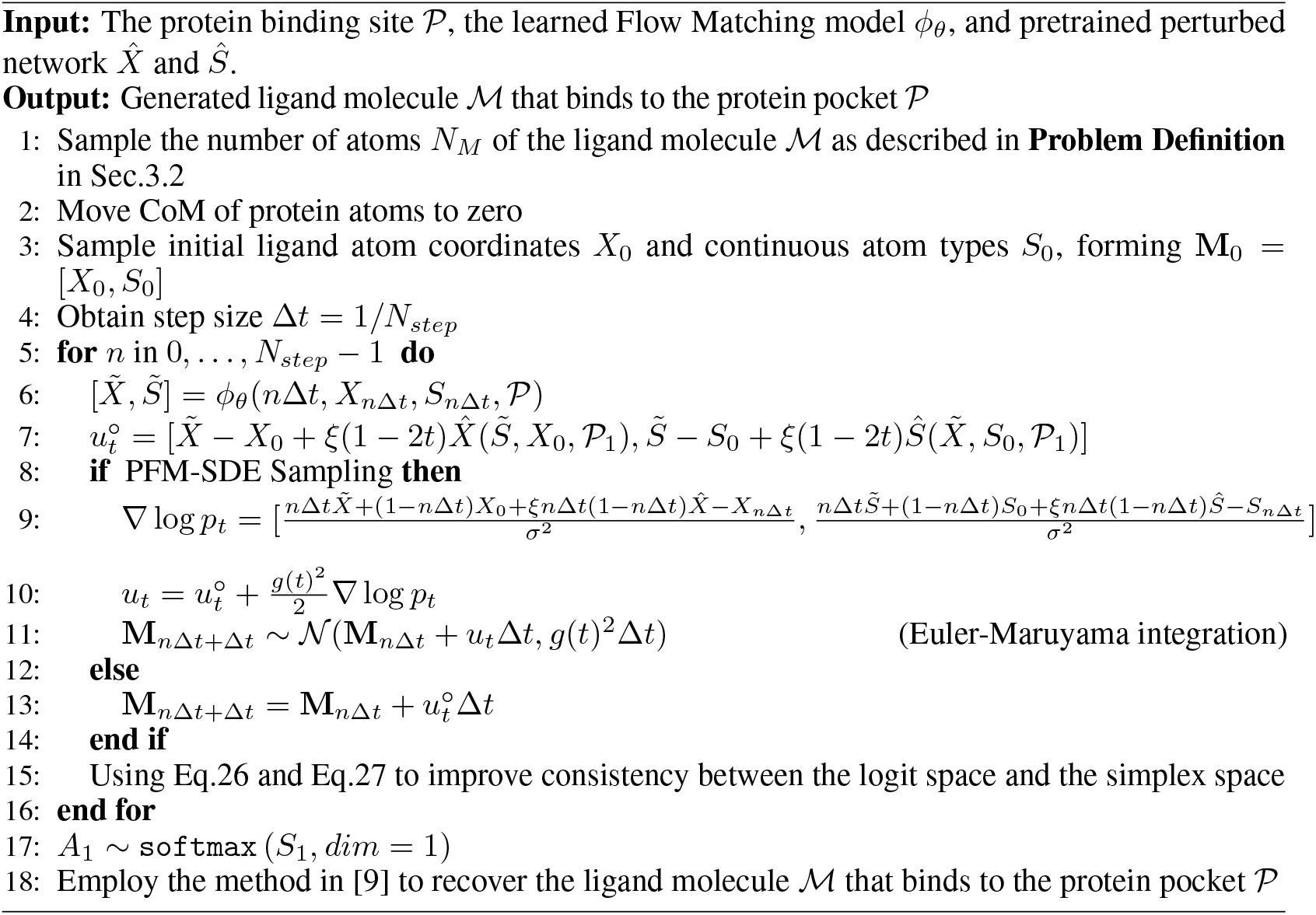

## C Implementation Details

### C.1 Initialization of Inputs

Following [9], protein atoms are represented using a combination of one-hot encoding for element types (H, C, N, O, S, Se) and amino acid types (20 types), while ligand atoms are encoded using one-hot representations of their element types (C, N, O, F, P, S, Cl). Furthermore, an additional dimension is introduced to distinguish between protein and ligand atoms. Two linear layers map atoms from different sources to 128-dimensional latent space, respectively. Edge connections of graphs are encoded using a 4-dimensional one-hot vector indicating the connection type (intra-protein, intra-ligand, protein-ligand, or ligand-protein). Edge distances are embedded via radial basis functions located at 20 centers between 0 Å and 10 Å.

### C.2 Architecture

An 8-layer SE(3)-equivariant neural network with a hidden dimensionality of 128 and 16 attention heads is used for dot product attention-based message passing. At each layer, three knn graphs (*G*_*P*_, *G*_*M*_ and *G*_*I*_) are dynamically constructed based on distances, where *k*_*P*_ = 24, *k*_*M*_ = 32 and *k*_*I*_ = 32. Message passing and feature updates are conducted over these graphs according to Sec.3.5.

### C.3 Training Details

The model is trained using the Adam optimizer with learning rate of 5e-4, betas = (0.95, 0.999), batch size of 16, and clipped gradient norm of 8. During the training process for 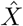 predictor and *Ŝ* predictor, Gaussian noise is added to *S*_1_ and *X*_1_ with standard deviations of 2.5 and 0.5, respectively, to adapt to vector field predictions with noise during inference. To ensure appropriate perturbation tendencies of 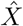, the operations described in Appendix A.3 are applied to *X*_0_ and *X*_1_. Throughout training, protein atom coordinates are augmented with Gaussian noise with a standard deviation of 0.1. We trained the parameterized PFM on a single NVIDIA GeForce RTX 4090 GPU, and it could converge within 62.5k steps.

### C.4 Hyperparameter Settings

In the experiments, Eq.6 uses *σ* = 0.1 and *ξ* = 1.5, while Eq.11 employs *γ* = 3.5 and uses 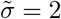. In Eq.12, *µ* = 0.75 and *v* = 10. For ablation studies, *ζ* = 2.53 in LPFM and RPFM to align with the peak values of perturbation terms in the PFM schedule. In Algorithm 2, *g*(*t*) is set to 0.1.

## D Additional Results

Tab.6 presents the training and inference times of PFM compared with two other diffusion-based models, with all models configured according to the settings specified in the respective paper. Training time refers to the duration required for model convergence, while inference time denotes the time taken to generate 10,000 valid molecules. All tests were conducted on the same NVIDIA GeForce RTX 4090 GPU.

Tab.7 highlights the positive effects of practices described in Appendix A.3 and Appendix A.1 on the model’s generative performance. PFM-True Loss (**PFM-TL**) indicates the use of 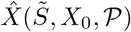 and 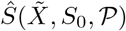 during training, while **PFM-IC** represents the use of independent coupling during the training process. *Training Loss* and *Test Loss* report the best loss during training and test, respectively.

Tab.8 presents an ablation study on the perturbation coefficient, showing that *ξ* = 1.5 is a generally better choice; *ξ* = 0 corresponds to BFM, while *ξ* = 1.5 corresponds to PFM.

**Table 8:** Ablation of perturbation coefficients.

| Methods | Vina Score (↓) | | Vina Min (↓) | | QED (↑) | | SA (↑) | |
| --- | --- | --- | --- | --- | --- | --- | --- | --- |
|  | Avg. | Med. | Avg. | Med. | Avg. | Med. | Avg. | Med. |
| $\xi = 0$ | -5.77 | -6.27 | -6.76 | -6.71 | 0.47 | 0.48 | 0.54 | 0.54 |
| $\xi = 0.5$ | -6.68 | -7.15 | -7.48 | -6.30 | 0.48 | 0.49 | 0.48 | 0.48 |
| $\xi = 1.0$ | -6.78 | -6.83 | -7.25 | -7.13 | 0.48 | 0.49 | 0.52 | 0.52 |
| $\xi = 1.5$ | -7.12 | -7.18 | -7.58 | -7.32 | 0.49 | 0.50 | 0.51 | 0.51 |

Fig.3 visualizes the influence of perturbation in PFM on molecular generation outcomes and processes. Specifically, **PFM (BFM sampling)** refers to molecule generation using only the vector field predictions from the BFM component, while **PFM (BFM part)** represents the BFM component’s contribution when generating molecules with the total PFM vector field predictions. **Molecule-level Comparison** of vector fields illustrate the average atomic contributions. The top row of Fig.3 shows performance improvements due to the perturbed conditional probability path, and the bottom two rows demonstrate how perturbations alter the vector fields. The vector field in the figure is not smooth, as it is produced by the trained network rather than the true marginal distribution. Fig.5 provides more visualizations.

**Figure 3:**
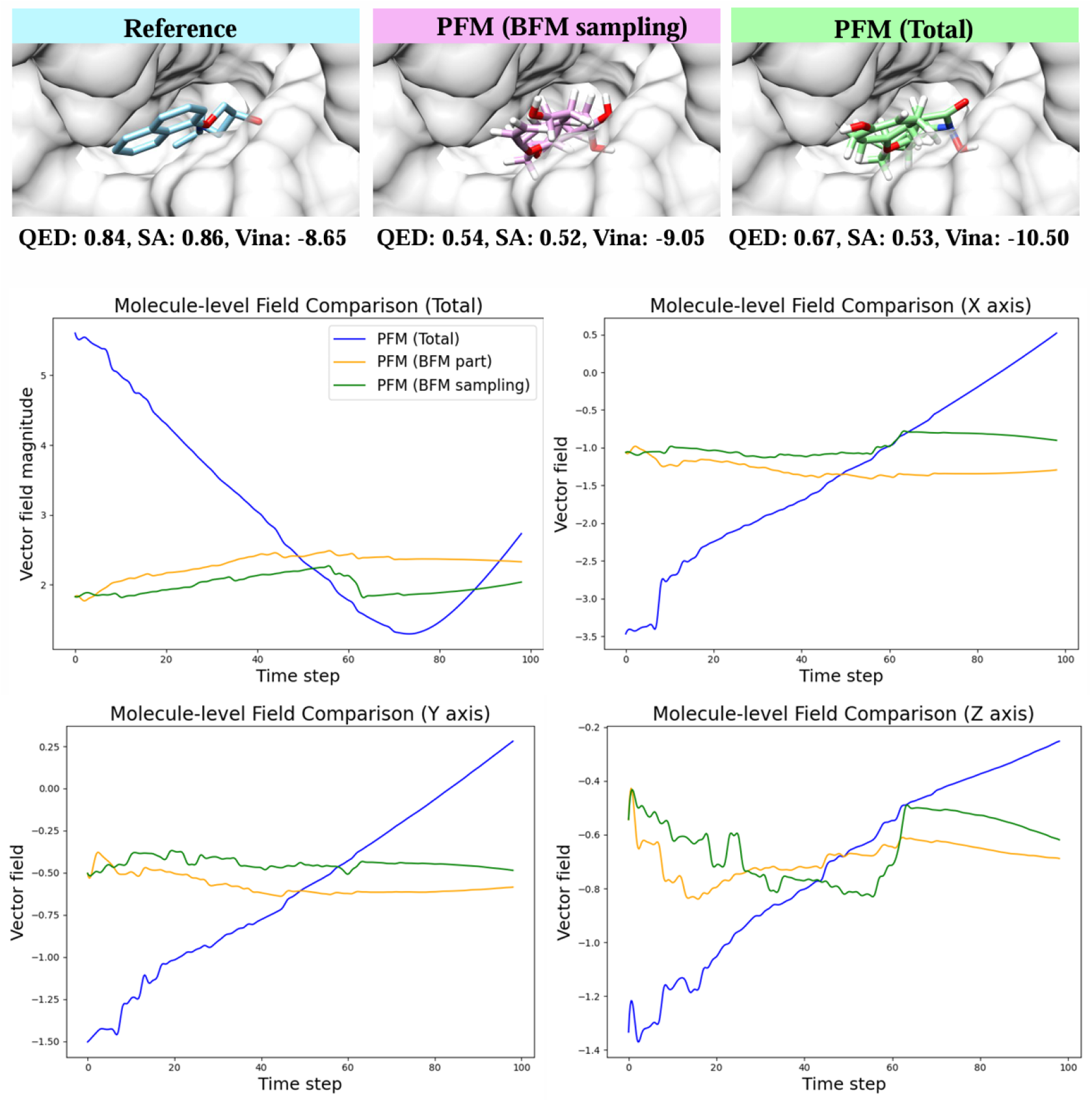
The influence of perturbation in PFM on molecular generation outcomes (top row) and processes (middle and bottom rows) on protein 2v3r.

Compared with TargetDiff, DecompDiff, and IPDiff, PFM achieves superior or competitive molecular generation results with fewer sampling steps (100 steps in the primary experiment). Fig.4 shows that further reducing the number of sampling steps does not lead to significant fluctuations in the quality of generated molecules. This indicates the method’s potential for faster molecule generation.

**Figure 4:**
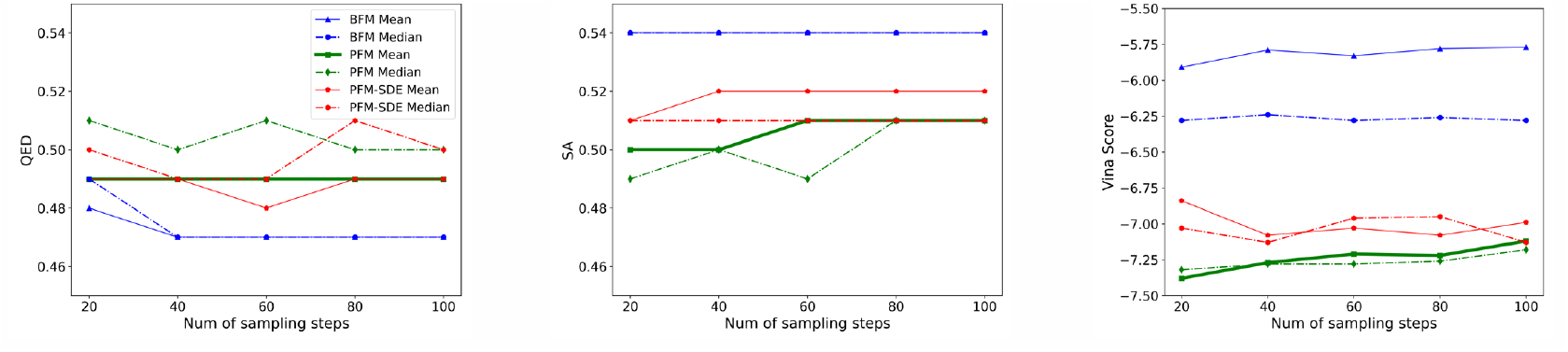
Ablation study on sampling step number. We compare PFM with BFM in terms of QED, SA, Vina Score under different sampling step number settings.

**Figure 5:**
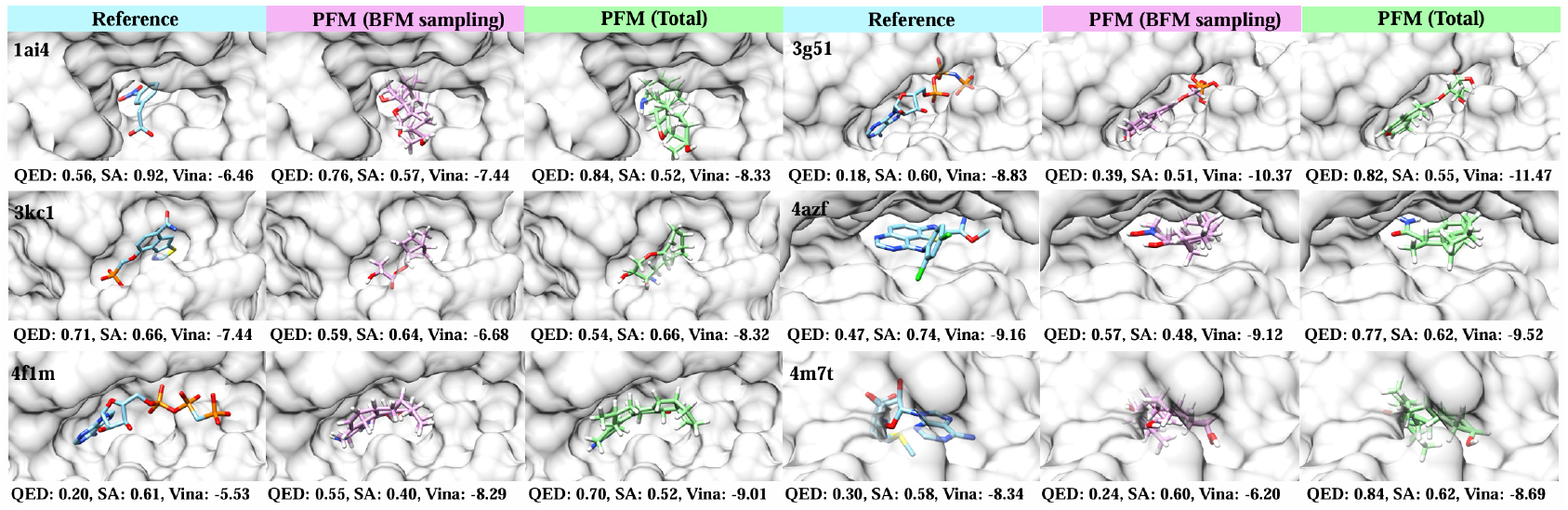
More examples of generated ligands for protein pockets. The generated molecules are produced by the trained PFM model. The results of PFM (BFM sampling) and PFM (Total) share the same noise initialization but differ in the vector field predictions used during the generation process.

In addition, we provide the visualization of more ligand molecules generated by PFM, comparing with both reference and IPDiff [11], as shown in Fig.6.

**Figure 6:**
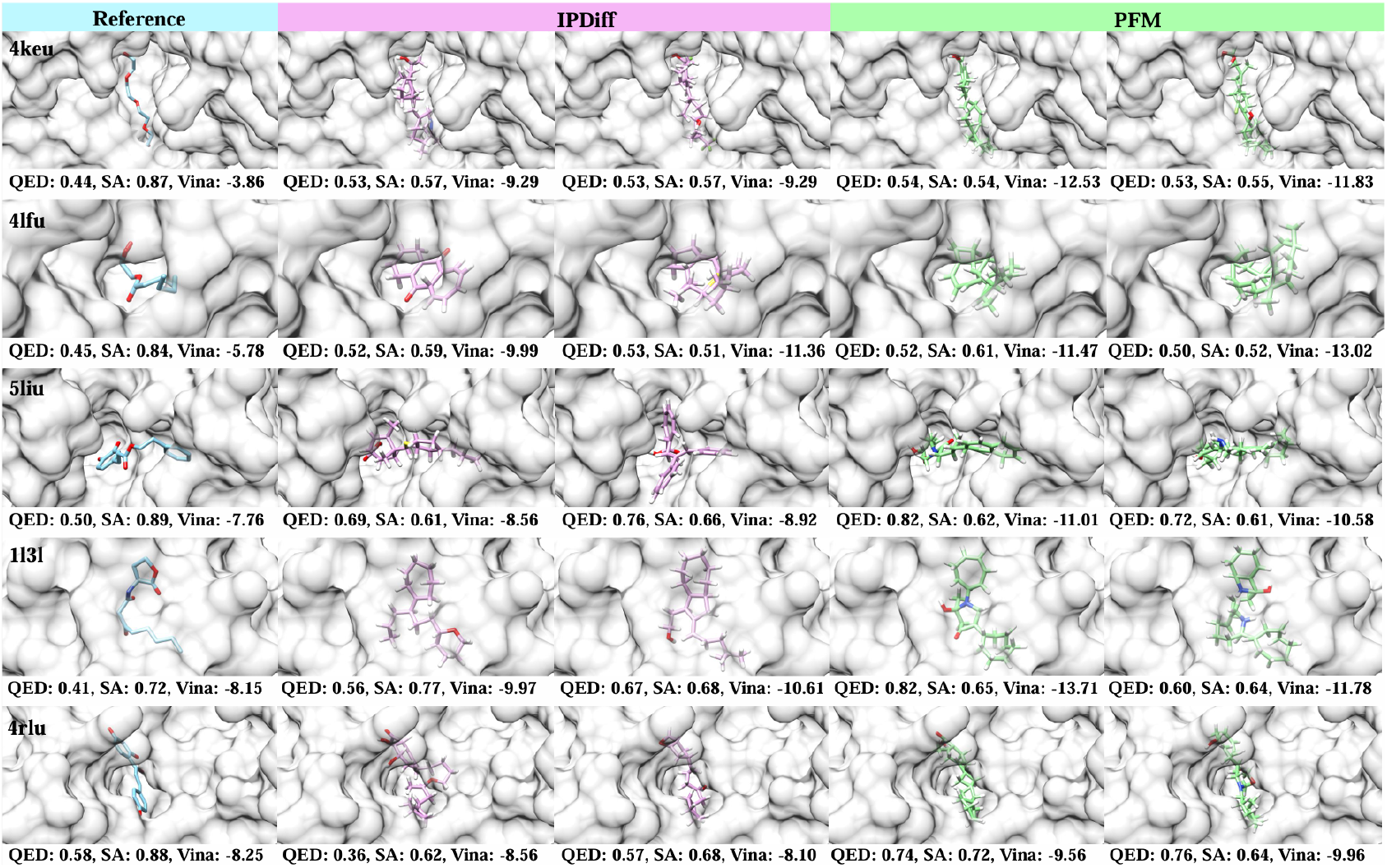
More examples of generated ligands for protein pockets. Carbon atoms in reference ligands, ligands generated by IPDiff [11] and PFM are visualized in blue, pink and green, respectively. QED, SA and Vina Score are reported for each molecule.

